## Supplementary File for "Revealing the drivers of antibiotic resistance trends in *Streptococcus pneumoniae* amidst the 2020 COVID-19 pandemic: Insights from mathematical modeling"

### Appendix 1

#### Literature search

We searched PubMed on December 4, 2023, for published mathematical modeling studies describing coinfection with SARS-CoV-2 and antibiotic-resistant bacteria and including epidemiological transmission of both pathogens. We used the following search query:

("model*"[Title/Abstract] OR "models, theoretical"[MeSH Terms] OR "simulat*"[Title/Abstract])

AND

("mathematic*"[MeSH Terms] OR "compartmental"[Title/Abstract] OR "agent-based"[Title/Abstract] OR "individual-based"[Title/Abstract] OR "stochastic*"[Title/Abstract] OR "deterministic"[Title/Abstract] OR "computer-generated"[Title/Abstract] OR "computer simulation"[MeSH Terms] OR "computer"[Title/Abstract] OR "computational"[Title/Abstract] OR "mathematical*"[Title/Abstract] OR "theoretical"[Title/Abstract] OR "prediction"[Title/Abstract] OR "simulat*"[Title/Abstract])

AND

("outbreak*"[Title/Abstract] OR "dynamic*"[Title/Abstract] OR "disseminat*"[Title/Abstract] OR "propagat*"[Title/Abstract] OR "transmi*"[Title/Abstract] OR "cross infection/transmission"[MeSH Terms] OR "spread*"[Title/Abstract] OR "diffus*"[Title/Abstract] OR "circulati*"[Title/Abstract] OR "disease activity"[Title/Abstract] OR "epidemiology"[Title/Abstract] OR "co-infection"[Title/Abstract])

AND

("COVID-19"[Title/Abstract] OR "SARS-CoV-2"[Title/Abstract])

AND

("bacteria*"[Title/Abstract] OR "antibiotic resist*"[Title/Abstract] OR "antibiotic-resist*"[Title/Abstract] OR "antimicrobial resist*"[Title/Abstract] OR "antimicrobial-resist*"[Title/Abstract]),

which returned 153 papers. One published study developed a mathematical model including the transmission of both SARS-CoV-2 and antibiotic-resistant bacteria but is specific to the hospital setting (Smith et al., 2023). Two bacterial transmission models implicitly accounted for COVID-19 by simulating pandemic-associated social distancing interventions/lockdowns, and corresponding reductions in (i) close-proximity contacts, in the context of transmission of antibiotic-resistance genes among human microbiota (Rebelo et al., 2021), and (ii) sexual contacts, in the context of transmission of HIV, gonorrhea and chlamydia (Jenness et al., 2021). A third model of *Neisseria meningitidis* transmission also evaluated colonization and infection dynamics over the pandemic period (Cascante-Vega et al., 2023). However, none of these models included SARS-CoV-2 infection and transmission, and only the first considered antibiotic resistance. One further transmission modelling study included coinfection with SARS-CoV-2 and influenza-like illness but did not consider bacteria (Bhowmick et al., 2023). Our search also returned some statistical models used to analyze epidemiological trends in bacterial coinfection subsequent to SARS-CoV-2 infection (Zhou et al., 2023) and changes in the incidence of bacterial disease during the pandemic (Shaw et al., 2023), but these did not use transmission modelling approaches to understand or quantify the underlying epidemiological mechanisms driving these trends. Several other modelling studies have described within-host molecular dynamics or airborne fluid particle dynamics, but do not describe pathogen transmission at the host population level. No other relevant articles were identified.

### Appendix 2

#### Model equations:

Compartment notation:

$X_j^i$

$X$ = SARS-CoV-2 status (S = susceptible, E = exposed, I = infected, and R = recovered)
$i$ = bacterial carriage (no superscript = no carriage, S = antibiotic-sensitive carriage, R = antibiotic-resistant carriage, SR = dual carriage, SS = dual antibiotic-sensitive carriage, RR = dual antibiotic-resistant carriage)
$j$ = antibiotic exposure (no subscript = no antibiotic exposure, $a$ = exposed to baseline antibiotic treatment, $az$ = exposed to azithromycin treatment)

Total population size:

$$N = S + S^S + S^R + S^{SR} + S^{SS} + S^{RR} + S_a + S_a^S + S_a^R + S_a^{SR} + S_a^{SS} + S_a^{RR} + E + E^S + E^R + E^{SR} + E^{SS} + E^{RR} + E_a + E_a^S + E_a^R + E_a^{SR} + E_a^{SS} + E_a^{RR} + I + I^S + I^R + I^{SR} + I^{SS} + I^{RR} + I_a + I_a^S + I_a^R + I_a^{SR} + I_a^{SS} + I_a^{RR} + I_{az} + I_{az}^S + I_{az}^R + I_{az}^{SR} + I_{az}^{SS} + I_{az}^{RR} + R + R^S + R^R + R^{SR} + R^{SS} + R^{RR} + R_a + R_a^S + R_a^R + R_a^{SR} + R_a^{SS} + R_a^{RR} + I_a^{RR} + R_{az} + R_{az}^S + R_{az}^R + R_{az}^{SR} + R_{az}^{SS} + R_{az}^{RR}$$

Forces of infection:

*S. pneumoniae*

$$\lambda_S = \beta_S\theta_\beta(S^S + 2qp_sS^{SR} + 2qS^{SS} + S_a^S + 2qp_sS_a^{SR} + 2qS_a^{SS} + E^S + 2qp_sE^{SR} + 2qE^{SS} + E_a^S + 2qp_sE_a^{SR} + 2qE_a^{SS} + I^S + 2qp_sI^{SR} + 2qI^{SS} + I_a^S + 2qp_sI_a^{SR} + 2qI_a^{SS} + I_{az}^S + 2qp_sI_{az}^{SR} + 2qI_{az}^{SS} + R^S + 2qp_sR^{SR} + 2qR^{SS} + R_a^S + 2qp_sR_a^{SR} + 2qR_a^{SS} + R_{az}^S + 2qp_sR_{az}^{SR} + 2qR_{az}^{SS})/N$$

$$\lambda_R = \beta_Sf\theta_\beta(S^R + 2q(1-p_s)S^{SR} + 2qS^{RR} + S_a^R + 2q(1-p_s)S_a^{SR} + 2qS_a^{RR} + E^R + 2q(1-p_s)E^{SR} + 2qE^{RR} + E_a^R + 2q(1-p_s)E_a^{SR} + 2qE_a^{RR} + I^R + 2q(1-p_s)I^{SR} + 2qI^{RR} + I_a^R + 2q(1-p_s)I_a^{SR} + 2qI_a^{RR} + I_{az}^R + 2q(1-p_s)I_{az}^{SR} + 2qI_{az}^{RR} + R^R + 2q(1-p_s)R^{SR} + 2qR^{RR} + R_a^R + 2q(1-p_s)R_a^{SR} + 2qR_a^{RR} + R_{az}^R + 2q(1-p_s)R_{az}^{SR} + 2qR_{az}^{RR})/N$$

$$\lambda_{XS} = \beta_S\theta_\beta \left(S^S + p_{single}(2qp_sS^{SR} + 2qS^{SS}) + S_a^S + p_{single}(2qp_sS_a^{SR} + 2qS_a^{SS}) + E^S + p_{single}(2qp_sE^{SR} + 2qE^{SS}) + E_a^S + p_{single}(2qp_sE_a^{SR} + 2qE_a^{SS}) + I^S + p_{single}(2qp_sI^{SR} + 2qI^{SS}) + I_a^S + p_{single}(2qp_sI_a^{SR} + 2qI_a^{SS}) + I_{az}^S + p_{single}(2qp_sI_{az}^{SR} + 2qI_{az}^{SS}) + R^S + p_{single}(2qp_sR^{SR} + 2qR^{SS}) + R_a^S + p_{single}(2qp_sR_a^{SR} + 2qR_a^{SS}) + R_{az}^S + p_{single}(2qp_sR_{az}^{SR} + 2qR_{az}^{SS})\right)/N$$

$$\lambda_{XR} = \beta_S f\theta_\beta \left(S^R + p_{single}(2q(1-p_s)S^{SR} + 2qS^{RR}) + S_a^R + p_{single}(2q(1-p_s)S_a^{SR} + 2qS_a^{RR}) + E^R + p_{single}(2q(1-p_s)E^{SR} + 2qE^{RR}) + E_a^R + p_{single}(2q(1-p_s)E_a^{SR} + 2qE_a^{RR}) + I^R + p_{single}(2q(1-p_s)I^{SR} + 2qI^{RR}) + I_a^R + p_{single}(2q(1-p_s)I_a^{SR} + 2qI_a^{RR}) + I_{az}^R + p_{single}(2q(1-p_s)I_{az}^{SR} + 2qI_{az}^{RR}) + R^R + p_{single}(2q(1-p_s)R^{SR} + 2qR^{RR}) + R_a^R + p_{single}(2q(1-p_s)R_a^{SR} + 2qR_a^{RR}) + R_{az}^R + p_{single}(2q(1-p_s)R_{az}^{SR} + 2qR_{az}^{RR})\right)/N$$

$$\lambda_{XSR} = \left((1-p_{single})\left(\beta_S\theta_\beta p_s + \beta_S\theta_\beta f(1-p_s)\right)2q(S^{SR} + S_a^{SR} + E^{SR} + E_a^{SR} + I^{SR} + I_a^{SR} + I_{az}^{SR} + R^{SR} + R_a^{SR} + R_{az}^{SR})\right)/N$$

$$\lambda_{XSS} = \left((1-p_{single})\beta_S\theta_\beta 2q(S^{SS} + S_a^{SS} + E^{SS} + E_a^{SS} + I^{SS} + I_a^{SS} + I_{az}^{SS} + R^{SS} + R_a^{SS} + R_{az}^{SS})\right)/N$$

$$\lambda_{XRR} = \left((1-p_{single})\beta_S\theta_\beta f2q(S^{RR} + S_a^{RR} + E^{RR} + E_a^{RR} + I^{RR} + I_a^{RR} + I_{az}^{RR} + R^{RR} + R_a^{RR} + R_{az}^{RR})\right)/N$$

SARS-CoV-2:

$$\lambda_C = \beta_C\theta_C((I + I^S + I^R + I^{SR} + I^{SS} + I^{RR} + I_a + I_a^S + I_a^R + I_a^{SR} + I_a^{SS} + I_a^{RR} + I_{az} + I_{az}^S + I_{az}^R + I_{az}^{SR} + I_{az}^{SS} + I_{az}^{RR})/N$$

Clearance rates of pneumococcal carriage:

$$\gamma^S = \gamma^R = \gamma^{SS} = \gamma^{RR} = \gamma^{SR}$$

The equations for SARS-CoV-2 susceptible individuals (no antibiotic treatment):

$$\frac{dS}{dt} = -(\lambda_{XS} + \lambda_{XR} + \lambda_{XSR} + \lambda_{XSS} + \lambda_{XRR})S + \gamma^S(S^S + S^{SS}) + \gamma^R(S^R + S^{SR} + S^{RR} + S_a^R + S_a^{RR}) + (\gamma^S + \omega)(S_a^S + S_a^{SS}) - \tau aS + rS_a - \lambda_C S$$

$$\frac{dS^S}{dt} = \lambda_{XS}S - \gamma^S S^S - k\lambda_S S^S - k\lambda_R S^S - \tau a S^S - \lambda_C S^S$$

$$\frac{dS^R}{dt} = \lambda_{XR}S - \gamma^R S^R - k\lambda_S S^R - k\lambda_R S^R - \tau a S^R + rS_a^R + (\gamma^S + \omega)S_a^{SR} - \lambda_C S^R$$

$$\frac{dS^{SR}}{dt} = \lambda_{XSR}S - \gamma^R S^{SR} + k\lambda_R S^S + k\lambda_S S^R - kc\lambda_S S^{SR} - kc\lambda_R S^{SR} + 2kc\lambda_R S^{SS} + 2kc\lambda_S S^{RR} - \tau a S^{SR} - \lambda_C S^{SR}$$

$$\frac{dS^{SS}}{dt} = \lambda_{XSS}S - \gamma^S S^{SS} + k\lambda_S S^S + kc\lambda_S S^{SR} - 2kc\lambda_R S^{SS} - \tau a S^{SS} - \lambda_C S^{SS}$$

$$\frac{dS^{RR}}{dt} = \lambda_{XRR}S - \gamma^R S^{RR} + k\lambda_R S^R + kc\lambda_R S^{SR} - 2kc\lambda_S S^{RR} - \tau a S^{RR} + rS_a^{RR} - \lambda_C S^{RR}$$

The equations for SARS-CoV-2 susceptible individuals (with antibiotic treatment):

$$\frac{dS_a}{dt} = -(\lambda_{XR} + \lambda_{XRR})S_a + \tau a S - rS_a - \lambda_C S_a$$

$$\frac{dS_a^S}{dt} = -(\gamma^S + \omega)S_a^S + \tau a S^S - k\lambda_R S_a^S - \lambda_C S_a^S$$

$$\frac{dS_a^R}{dt} = -\gamma^R S_a^R + \lambda_{XR} S_a + \tau a S^R - rS_a^R - \lambda_C S_a^R$$

$$\frac{dS_a^{SR}}{dt} = -(\gamma^S + \omega)S_a^{SR} + k\lambda_R S_a^S + 2kc\lambda_R S_a^{SS} + \tau a S^{SR} - \lambda_C S_a^{SR}$$

$$\frac{dS_a^{SS}}{dt} = -(\gamma^S + \omega)S_a^{SS} - 2kc\lambda_R S_a^{SS} + \tau a S^{SS} - \lambda_C S_a^{SS}$$

$$\frac{dS_a^{RR}}{dt} = \lambda_{XRR} S_a + \tau a S^{RR} - rS_a^{RR} - \gamma^R S_a^{RR} - \lambda_C S_a^{RR}$$

The equations for SARS-CoV-2 exposed individuals (no antibiotic treatment):

$$\frac{dE}{dt} = -(\lambda_{XS} + \lambda_{XR} + \lambda_{XSR} + \lambda_{XSS} + \lambda_{XRR})E + \gamma^S(E^S + E^{SS}) + \gamma^R(E^R + E^{SR} + E^{RR} + E_a^R + E_a^{RR}) + (\gamma^S + \omega)(E_a^S + E_a^{SS}) - \tau a E + rE_a + \lambda_C S - \varepsilon E$$

$$\frac{dE^S}{dt} = \lambda_{XS}E - \gamma^S E^S - k\lambda_S E^S - k\lambda_R E^S - \tau a E^S + \lambda_C S^S - \varepsilon E^S$$

$$\frac{dE^R}{dt} = \lambda_{XR}E - \gamma^R E^R - k\lambda_S E^R - k\lambda_R E^R - \tau a E^R + rE_a^R + \lambda_C S^R - \varepsilon E^R + (\gamma^S + \omega)E_a^{SR}$$

$$\frac{dE^{SR}}{dt} = \lambda_{XSR}E - \gamma^R E^{SR} + k\lambda_R E^S + k\lambda_S E^R - kc\lambda_S E^{SR} - kc\lambda_R E^{SR} + 2kc\lambda_R E^{SS} + 2kc\lambda_S E^{RR} - \tau a E^{SR} + \lambda_C S^{SR} - \varepsilon E^{SR}$$

$$\frac{dE^{SS}}{dt} = \lambda_{XSS}E - \gamma^S E^{SS} + k\lambda_S E^S + kc\lambda_S E^{SR} - 2kc\lambda_R E^{SS} - \tau a E^{SS} + \lambda_C S^{SS} - \varepsilon E^{SS}$$

$$\frac{dE^{RR}}{dt} = \lambda_{XRR}E - \gamma^R E^{RR} + k\lambda_R E^R + kc\lambda_R E^{SR} - 2kc\lambda_S E^{RR} - \tau a E^{RR} + rE_a^{RR} + \lambda_C S^{RR} - \varepsilon E^{RR}$$

The equations for SARS-CoV-2 exposed individuals (with antibiotic treatment):

$$\frac{dE_a}{dt} = -(\lambda_{XR} + \lambda_{XRR})E_a + \tau a E - rE_a + \lambda_C S_a - \varepsilon E_a$$

$$\frac{dE_a^S}{dt} = -(\gamma^S + \omega)E_a^S + \tau a E^S - k\lambda_R E_a^S + \lambda_C S_a^S - \varepsilon E_a^S$$

$$\frac{dE_a^R}{dt} = -\gamma^R E_a^R + \lambda_{XR}E_a + \tau a E^R - rE_a^R + \lambda_C E_a^R - \varepsilon E_a^R$$

$$\frac{dE_a^{SR}}{dt} = -(\gamma^S + \omega)E_a^{SR} + k\lambda_R E_a^S + 2kc\lambda_R E_a^{SS} + \tau a E^{SR} + \lambda_C S_a^{SR} - \varepsilon E_a^{SR}$$

$$\frac{dE_a^{SS}}{dt} = -(\gamma^S + \omega)E_a^{SS} - 2kc\lambda_R E_a^{SS} + \tau a E^{SS} + \lambda_C S_a^{SS} - \varepsilon E_a^{SS}$$

$$\frac{dE_a^{RR}}{dt} = \lambda_{XRR}E_a + \tau a E^{RR} - rE_a^{RR} - \gamma^R E_a^{RR} + \lambda_C S_a^{RR} - \varepsilon E_a^{RR}$$

The equations for SARS-CoV-2 infected individuals (no antibiotic treatment):

$$\frac{dI}{dt} = -(\lambda_{XS} + \lambda_{XR} + \lambda_{XSR} + \lambda_{XSS} + \lambda_{XRR})I + \gamma^S(I^S + I^{SS}) + \gamma^R(I^R + I^{SR} + I^{RR} + I_a^R + I_a^{RR}) + (\gamma^S + \omega)(I_a^S + I_a^{SS}) - (1 - p_{az})\tau a I + rI_a + \varepsilon E - \gamma^C I - p_{az}I + \gamma^R(I_{az}^R + I_{az}^{RR})$$

$$\frac{dI^S}{dt} = \lambda_{XS}I - \gamma^S I^S - k\lambda_S I^S - k\lambda_R I^S - (1 - p_{az})\tau a I^S + \varepsilon E^S - \gamma^C I^S - p_{az}I^S$$

$$\frac{dI^R}{dt} = \lambda_{XR}I - \gamma^R I^R - k\lambda_S I^R - k\lambda_R I^R + (\gamma^S + \omega)I_a^{SR} - (1 - p_{az})\tau a I^R + rI_a^R + \varepsilon E^R - \gamma^C I^R - p_{az}I^R$$

$$\frac{dI^{SR}}{dt} = \lambda_{XSR}I - \gamma^R I^{SR} + k\lambda_R I^S + k\lambda_S I^R - kc\lambda_S I^{SR} - kc\lambda_R I^{SR} + 2kc\lambda_R I^{SS} + 2kc\lambda_S I^{RR} - (1 - p_{az})\tau a I^{SR} + \varepsilon E^{SR} - \gamma^C I^{SR} - p_{az}I^{SR}$$

$$\frac{dI^{SS}}{dt} = \lambda_{XSS}I - \gamma^S I^{SS} + k\lambda_S I^S + kc\lambda_S I^{SR} - 2kc\lambda_R I^{SS} - (1 - p_{az})\tau a I^{SS} + \varepsilon E^{SS} - \gamma^C I^{SS} - p_{az}I^{SS}$$

$$\frac{dI^{RR}}{dt} = \lambda_{XRR}I - \gamma^R I^{RR} + k\lambda_R I^R + kc\lambda_R I^{SR} - 2kc\lambda_S I^{RR} - (1 - p_{az})\tau a I^{RR} + rI_a^{RR} + \varepsilon E^{RR} - \gamma^C I^{RR} - p_{az}I^{RR}$$

The equations for SARS-CoV-2 infected individuals (with antibiotic treatment):

$$\frac{dI_a}{dt} = -(\lambda_{XR} + \lambda_{XRR})I_a + (1 - p_{az})\tau a I - rI_a + \varepsilon E_a - \gamma^C I_a$$

$$\frac{dI_a^S}{dt} = -(\gamma^S + \omega)I_a^S + (1 - p_{az})\tau a I^S - k\lambda_R I_a^S + \varepsilon E_a^S - \gamma^C I_a^S$$

$$\frac{dI_a^R}{dt} = -\gamma^R I_a^R + \lambda_{XR}I_a + (1 - p_{az})\tau a I^R - rI_a^R + \varepsilon E_a^R - \gamma^C I_a^R$$

$$\frac{dI_a^{SR}}{dt} = -(\gamma^S + \omega)I_a^{SR} + k\lambda_R I_a^S + 2kc\lambda_R I_a^{SS} + (1 - p_{az})\tau a I^{SR} + \varepsilon E_a^{SR} - \gamma^C I_a^{SR}$$

$$\frac{dI_a^{SS}}{dt} = -(\gamma^S + \omega)I_a^{SS} - 2kc\lambda_R I_a^{SS} + (1 - p_{az})\tau a I^{SS} + \varepsilon E_a^{SS} - \gamma^C I_a^{SS}$$

$$\frac{dI_a^{RR}}{dt} = \lambda_{XRR}I_a + (1 - p_{az})\tau a I^{RR} - rI_a^{RR} - \gamma^R I_a^{RR} + \varepsilon E_a^{RR} - \gamma^C I_a^{RR}$$

The equations for SARS-CoV-2 infected individuals (with azithromycin treatment):

$$\frac{dI_{az}}{dt} = p_{az}I - \gamma^C I_{az} - (\lambda_{XR} + \lambda_{XRR})I_{az} + (\gamma^S + \omega)(I_{az}^S + I_{az}^{SS})$$

$$\frac{dI_{az}^S}{dt} = p_{az}I^S - \gamma^C I_{az}^S - (\gamma^S + \omega)I_{az}^S - k\lambda_R I_{az}^S$$

**Appendix 2**

$$\frac{dI_{az}^{R}}{dt} = p_{az}I^{R} - \gamma^{C}I_{az}^{R} + \lambda_{XR}I_{az} - \gamma^{R}I_{az}^{R} + (\gamma^{S} + \omega)I_{az}^{SR}$$

$$\frac{dI_{az}^{SR}}{dt} = p_{az}I^{SR} - \gamma^{C}I_{az}^{SR} - (\gamma^{S} + \omega)I_{az}^{SR} + k\lambda_{R}I_{az}^{S} + 2kc\lambda_{R}I_{az}^{SS}$$

$$\frac{dI_{az}^{SS}}{dt} = p_{az}I^{SS} - \gamma^{C}I_{az}^{SS} - (\gamma^{S} + \omega)I_{az}^{SS} - 2kc\lambda_{R}I_{az}^{SS}$$

$$\frac{dI_{az}^{RR}}{dt} = p_{az}I^{RR} - \gamma^{C}I_{az}^{RR} + \lambda_{XRR}I_{az} - \gamma^{R}I_{az}^{RR}$$

The equations for SARS-CoV-2 recovered individuals (no antibiotic treatment):

$$\begin{aligned}\frac{dR}{dt} = -(\lambda_{XS} + \lambda_{XR} + \lambda_{XSR} + \lambda_{XSS} + \lambda_{XRR})R + \gamma^{S}(R^{S} + R^{SS}) \\ + \gamma^{R}(R^{R} + R^{SR} + R^{RR} + R_{a}^{R} + R_{a}^{RR}) + (\gamma^{S} + \omega)(R_{a}^{S} + R_{a}^{SS}) - \tau aR + rR_{a} \\ + \gamma^{C}I + r_{az}R_{az} + \gamma^{R}(R_{az}^{R} + R_{az}^{RR})\end{aligned}$$

$$\frac{dR^{S}}{dt} = \lambda_{XS}R - \gamma^{S}R^{S} - k\lambda_{S}R^{S} - k\lambda_{R}R^{S} - \tau aR^{S} + \gamma^{C}I^{S} + r_{az}R_{az}^{S}$$

$$\frac{dR^{R}}{dt} = \lambda_{XR}R - \gamma^{R}R^{R} - k\lambda_{S}R^{R} - k\lambda_{R}R^{R} + (\gamma^{S} + \omega)R_{a}^{SR} - \tau aR^{R} + rR_{a}^{R} + \gamma^{C}I^{R} + r_{az}R_{az}^{R}$$

$$\begin{aligned}\frac{dR^{SR}}{dt} = \lambda_{XSR}R - \gamma^{R}R^{SR} + k\lambda_{R}R^{S} + k\lambda_{S}R^{R} - kc\lambda_{S}R^{SR} - kc\lambda_{R}R^{SR} + 2kc\lambda_{R}R^{SS} + 2kc\lambda_{S}R^{RR} \\ - \tau aR^{SR} + \gamma^{C}I^{SR} + r_{az}R_{az}^{SR}\end{aligned}$$

$$\frac{dR^{SS}}{dt} = \lambda_{XSS}R - \gamma^{S}R^{SS} + k\lambda_{S}R^{S} + kc\lambda_{S}R^{SR} - 2kc\lambda_{R}R^{SS} - \tau aR^{SS} + \gamma^{C}I^{SS} + r_{az}R_{az}^{SS}$$

$$\begin{aligned}\frac{dR^{RR}}{dt} = \lambda_{XRR}R - \gamma^{R}R^{RR} + k\lambda_{R}R^{R} + kc\lambda_{R}R^{SR} - 2kc\lambda_{S}R^{RR} - \tau aR^{RR} + rR_{a}^{RR} + \gamma^{C}I^{RR} \\ + r_{az}R_{az}^{RR}\end{aligned}$$

The equations for SARS-CoV-2 recovered individuals (with antibiotic treatment):

$$\frac{dR_{a}}{dt} = \gamma^{C}I_{a} + \tau aR - rR_{a} - \lambda_{XR}R_{a} - \lambda_{XRR}R_{a}$$

$$\frac{dR_a^S}{dt} = \gamma^C I_a^S + \tau a R^S - (\gamma^S + \omega) R_a^S - k\lambda_R R_a^S$$

$$\frac{dR_a^R}{dt} = \gamma^C I_a^R + \tau a R^R - r R_a^R + \lambda_{XR} R_a - \gamma^R R_a^R$$

$$\frac{dR_a^{SR}}{dt} = \gamma^C I_a^{SR} + \tau a R^{SR} - (\gamma^S + \omega) R_a^{SR} + k\lambda_R R_a^S + 2kc\lambda_R R_a^{SS}$$

$$\frac{dR_a^{SS}}{dt} = \gamma^C I_a^{SS} + \tau a R^{SS} - (\gamma^S + \omega) R_a^{SS} - 2kc\lambda_R R_a^{SS}$$

$$\frac{dR_a^{RR}}{dt} = \gamma^C I_a^{RR} + \tau a R^{RR} - r R_a^{RR} + \lambda_{XRR} R_a - \gamma^R R_a^{RR}$$

The equations for SARS-CoV-2 recovered individuals (with azithromycin treatment):

$$\frac{dR_{az}}{dt} = \gamma^C I_{az} - r_{az} R_{az} - \lambda_{XR} R_{az} - \lambda_{XRR} R_{az} + (\gamma^S + \omega)(R_{az}^S + R_{az}^{SS})$$

$$\frac{dR_{az}^S}{dt} = \gamma^C I_{az}^S - r_{az} R_{az}^S - (\gamma^S + \omega) R_{az}^S - k\lambda_R R_{az}^S$$

$$\frac{dR_{az}^R}{dt} = \gamma^C I_{az}^R - r_{az} R_{az}^R + \lambda_{XR} R_{az} - \gamma^R R_{az}^R + (\gamma^S + \omega) R_{az}^{SR}$$

$$\frac{dR_{az}^{SR}}{dt} = \gamma^C I_{az}^{SR} - r_{az} R_{az}^{SR} - (\gamma^S + \omega) R_{az}^{SR} + k\lambda_R R_{az}^S + 2kc\lambda_R R_{az}^{SS}$$

$$\frac{dR_{az}^{SS}}{dt} = \gamma^C I_{az}^{SS} - r_{az} R_{az}^{SS} - (\gamma^S + \omega) R_{az}^{SS} - 2kc\lambda_R R_{az}^{SS}$$

$$\frac{dR_{az}^{RR}}{dt} = \gamma^C I_{az}^{RR} - r_{az} R_{az}^{RR} + \tau a R^{RR} - r R_a^{RR} + \lambda_{XRR} R_a - \gamma^R R_a^{RR}$$

The equations for invasive pneumococcal disease (IPD) incidence caused by sensitive and resistant strains:

$$\frac{dIPD^S}{dt} = p_{inv} IPD_{risk} \left( S^S + S^{SS} + \frac{1}{2} S^{SR} + E^S + E^{SS} + \frac{1}{2} E^{SR} + I^S + I^{SS} + \frac{1}{2} I^{SR} + R^S + R^{SS} + \frac{1}{2} R^{SR} \right)$$

**Appendix 2**

$$\frac{dIPD^R}{dt} = p_{inv}IPD_{risk}\Big(S^R + S^{RR} + \frac{1}{2}S^{SR} + E^R + E^{RR} + \frac{1}{2}E^{SR} + I^R + I^{RR} + \frac{1}{2}I^{SR} + R^R + R^{RR} + \frac{1}{2}R^{SR} + S_a^R + S_a^{RR} + S_a^{SR} + E_a^R + E_a^{RR} + E_a^{SR} + I_a^R + I_a^{RR} + I_a^{SR} + R_a^R + R_a^{RR} + R_a^{SR} + I_{az}^R + I_{az}^{RR} + I_{az}^{SR} + R_{az}^R + R_{az}^{RR} + R_{az}^{SR}\Big)$$

#### Equations for the prevalence of pneumococcal carriage in the population:

Prevalence of antibiotic-sensitive pneumococcal carriage is calculated as:

$$S_{prop} = \Big(S^S + \frac{1}{2}S^{SR} + S^{SS} + S_a^S + \frac{1}{2}S_a^{SR} + S_a^{SS} + E^S + \frac{1}{2}E^{SR} + E^{SS} + E_a^S + \frac{1}{2}E_a^{SR} + E_a^{SS} + I^S + \frac{1}{2}I^{SR} + I^{SS} + I_a^S + \frac{1}{2}I_a^{SR} + I_a^{SS} + I_{az}^S + \frac{1}{2}I_{az}^{SR} + I_{az}^{SS} + R^S + \frac{1}{2}R^{SR} + R^{SS} + R_a^S + \frac{1}{2}R_a^{SR} + R_a^{SS} + R_{az}^S + \frac{1}{2}R_{az}^{SR} + R_{az}^{SS}\Big)/Total\ carriage$$

Prevalence of antibiotic-resistant pneumococcal carriage is calculated as:

$$R_{prop} = \Big(S^R + \frac{1}{2}S^{SR} + S^{RR} + S_a^R + \frac{1}{2}S_a^{SR} + S_a^{RR} + E^R + \frac{1}{2}E^{SR} + E^{RR} + E_a^R + \frac{1}{2}E_a^{SR} + E_a^{RR} + I^R + \frac{1}{2}I^{SR} + I^{RR} + I_a^R + \frac{1}{2}I_a^{SR} + I_a^{RR} + I_{az}^R + \frac{1}{2}I_{az}^{SR} + I_{az}^{RR} + R^R + \frac{1}{2}R^{SR} + R^{RR} + R_a^R + \frac{1}{2}R_a^{SR} + R_a^{RR} + R_{az}^R + \frac{1}{2}R_{az}^{SR} + R_{az}^{RR}\Big)/Total\ carriage$$

Total pneumococcal carriage is calculated as:

$$Total\ carriage = S_{prop} + R_{prop}$$

Prevalence of total pneumococcal carriage is calculated as:

$$Total_{prop} = Total\ carriage/N$$

#### Equations for within-host interactions:

Invasive pneumococcal disease (IPD) incidence implementing within-host interactions is only accounted for in the equations for IPD incidence outside of the model equations, and hence has no impact on model dynamics.

Daily IPD incidence, for the antibiotic-resistant strain is calculated as:

$$\frac{dIPD^R}{dt} = p_{inv}IPD_{risk}\Big(S^R + S^{RR} + \frac{1}{2}S^{SR} + E^R + E^{RR} + \frac{1}{2}E^{SR} + \boldsymbol{\psi_c}(I^R + I^{RR} + \frac{1}{2}I^{SR}) + R^R + R^{RR} + \frac{1}{2}R^{SR} + S_a^R + S_a^{RR} + S_a^{SR} + E_a^R + E_a^{RR} + E_a^{SR} + \boldsymbol{\psi_c}(I_a^R + I_a^{RR} + I_a^{SR}) + R_a^R + R_a^{RR} + R_a^{SR} + \boldsymbol{\psi_c}(I_{az}^R + I_{az}^{RR} + I_{az}^{SR}) + R_{az}^R + R_{az}^{RR} + R_{az}^{SR}\Big)$$

where $p_{inv}$ represents the bacterial invasion rate or progression of carriage to invasive disease and $\psi_c$ represents the co-infection interaction term that increases the rate of progression to invasive disease among colonized individuals who are also infected with SARS-CoV-2.

Daily IPD incidence for the antibiotic-sensitive strain is calculated as:

$$\frac{dIPD^S}{dt} = p_{inv}IPD_{risk}\Big(S^R + S^{RR} + \frac{1}{2}S^{SR} + E^R + E^{RR} + \frac{1}{2}E^{SR} + \boldsymbol{\psi_c}(I^R + I^{RR} + \frac{1}{2}I^{SR}) + R^R + R^{RR} + \frac{1}{2}R^{SR} + S_a^R + S_a^{RR} + S_a^{SR} + E_a^R + E_a^{RR} + E_a^{SR} + \boldsymbol{\psi_c}(I_a^R + I_a^{RR} + I_a^{SR}) + R_a^R + R_a^{RR} + R_a^{SR} + \boldsymbol{\psi_c}(I_{az}^R + I_{az}^{RR} + I_{az}^{SR}) + R_{az}^R + R_{az}^{RR} + R_{az}^{SR}\Big)$$

Due to large uncertainty in the values of $\psi_c$ we considered broad range in our analysis estimated from prior modelling studies (Domenech De Cellès et al., 2019; Opatowski et al., 2013). In the context of ecological interactions between *S. pneumoniae* and influenza, estimations show that influenza co-infection increases the rate of progression from *S. pneumonia* colonization to invasive pneumococcal disease between 49 and 146-fold, on average, depending on the age group (Domenech De Cellès et al., 2019; Opatowski et al., 2013). Therefore, we implemented the range $1 < \psi_c < 100$ for SARS-CoV-2.

#### Appendix 2 – Figures

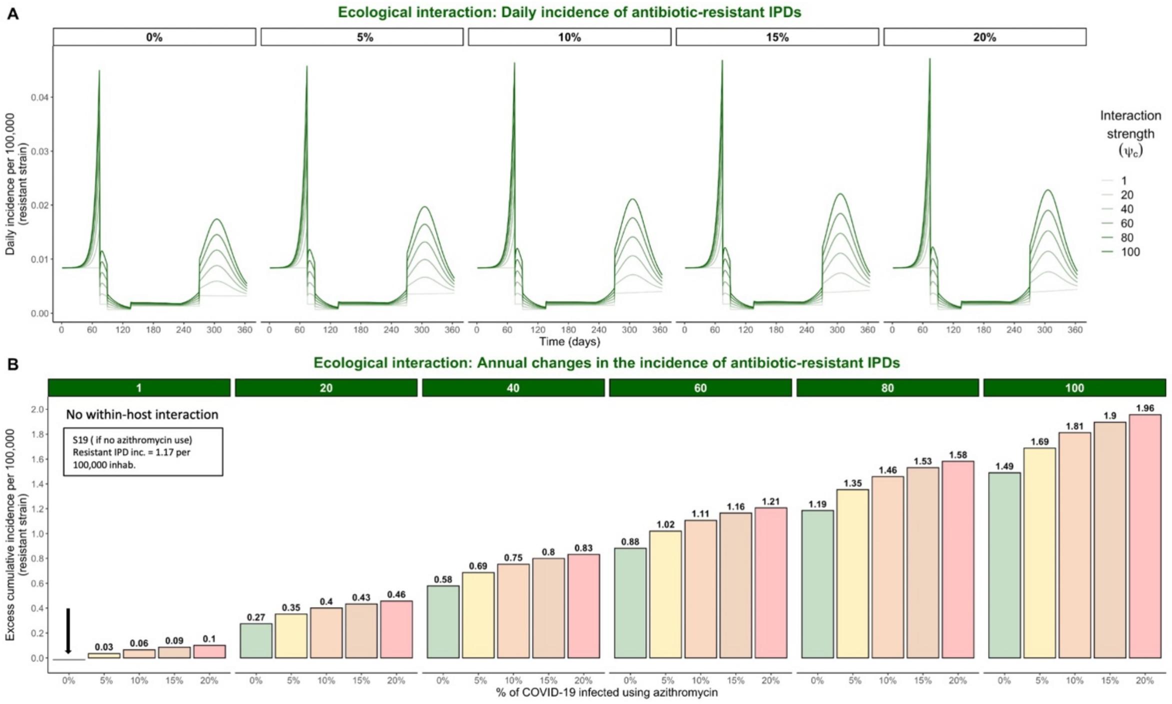


**Appendix 2 - Figure 1. Hypothetical within-host interactions in scenario S19 promote the incidence of antibiotic-resistant invasive pneumococcal disease (IPD). (A) Impacts of ecological interactions between pathogens.** When SARS-CoV-2 infection leads to faster progression from pneumococcal colonization to disease ($\psi_c > 1$), surges in COVID-19 cases accompanied by increasing levels of azithromycin use lead to substantial increases in the daily incidence of antibiotic-resistant IPD. **(B) Annual excess in cumulative IPD incidence due to synergistic within-host ecological interactions.** A rate of disease progression increased by a factor $\psi_c = 1$ (no within-host interaction) and $\psi_c = 40$ in scenario S19 applied to the general population resulted in approximately 0.06 and 0.75 additional cases of antibiotic-resistant disease per 100,000 inhabitants over the course of one year, respectively, compared to the scenario S19 assuming no within-host interaction and no azithromycin use indicated by the black arrow (1.17 cases/100,000 inhabitants).

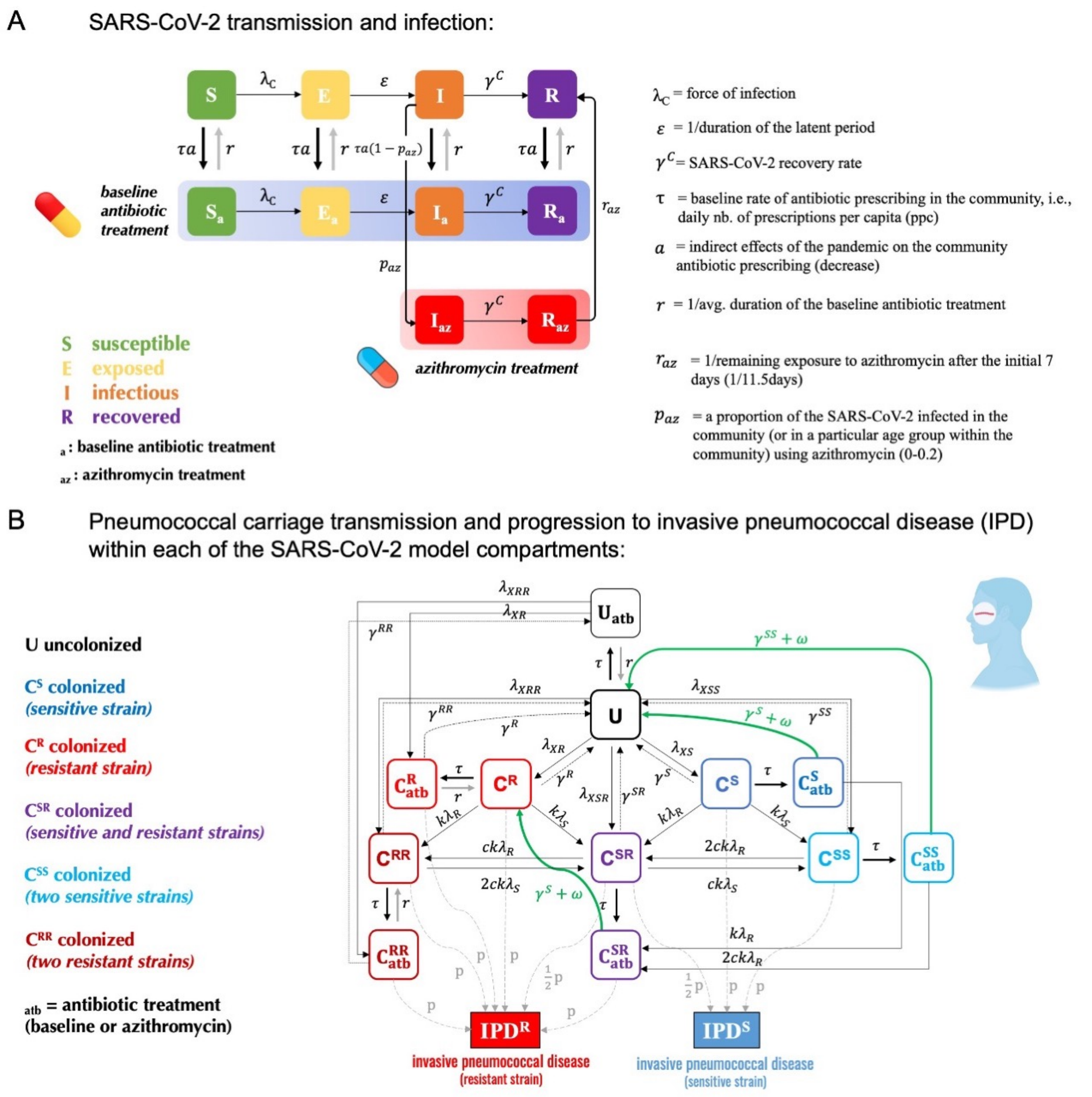


**Appendix 2 - Figure 2. Model schematic describing co-circulation of SARS-CoV-2 infection and pneumococcal carriage transmission. A.** Individuals in each of the SARS-CoV-2 transmission model compartments (S, E, I, R) can be either **B.** uncolonized (U) or colonized by the bacteria ($C^S$, $C^R$, $C^{SS}$, $C^{RR}$, $C^{SR}$) with or without concomitant antibiotic treatment (subscript $_{atb}$ denotes compartments exposed to antibiotics – both standard baseline antibiotic exposure and azithromycin exposure). Full list of model parameters can be found in Appendix 2 – Table 2.

#### TABLES

**Table 1.** Total number of invasive *S. pneumoniae* isolates from blood or cerebrospinal fluid tested in 2019 and 2020 and reported to EARS-Net (European Antimicrobial Resistance Surveillance Network).

| | *No. of invasive Streptococcus pneumoniae isolates in the EU/EEA* | | |
|---|---|---|---|
| **Country** | **2019** | **2020** | **% decrease in the total number of reported invasive isolates from 2019 to 2020** |
| *Austria* | 550 | 301 | 45.3% |
| *Belgium* | 1548 | 858 | 44.6% |
| *Bulgaria* | 46 | 28 | 39.1% |
| *Croatia* | 156 | 55 | 64.7% |
| *Czechia* | 387 | 204 | 47.3% |
| *Denmark* | 601 | 351 | 41.6% |
| *Estonia* | 161 | 80 | 50.3% |
| *Finland* | 678 | 293 | 56.8% |
| *France* | 1264 | 668 | 47.2% |
| *Germany* | 2035 | 1314 | 35.4% |
| *Hungary* | 222 | 124 | 44.1% |
| *Iceland* | 44 | 20 | 54.5% |
| *Ireland* | 348 | 136 | 60.9% |
| *Italy* | 1351 | 685 | 49.3% |
| *Latvia* | 79 | 42 | 46.8% |
| *Lithuania* | 120 | 96 | 20.0% |
| *Luxembourg* | 38 | 24 | 36.8% |
| *Malta* | 27 | 16 | 40.7% |
| *Netherlands* | 1552 | 997 | 35.8% |
| *Norway* | 507 | 243 | 52.1% |
| *Poland* | 364 | 165 | 54.7% |
| *Portugal* | 983 | 588 | 40.2% |
| *Romania* | 107 | 42 | 60.7% |
| *Slovakia* | 40 | 15 | 62.5% |
| *Slovenia* | 283 | 172 | 39.2% |
| *Spain* | 1038 | 611 | 41.1% |
| *Sweden* | 1071 | 551 | 48.6% |
| *United Kingdom* | 3468 | 1412 | 59.3% |

**Appendix 2**

**Table 2.** Summary of the model parameters:

| Symbol | Interpretation | Value(s) | References |
|---|---|---|---|
| **SARS-CoV-2 infection parameters:** | | | |
| $\boldsymbol{\beta_C}$ | transmission rate of SARS-CoV-2 | 0.46 $days^{-1}$ | (Liu et al., 2020) |
| $\boldsymbol{\gamma^C}$ | SARS-CoV-2 recovery rate (mild cases) | 1/7 $days^{-1}$ | (Lauer et al., 2020; Rhee et al., 2020) |
| $\boldsymbol{\varepsilon}$ | SARS-CoV-2 incubation rate (latent period from exposed to infectious state) | 1/5 $days^{-1}$ | (Elias et al., 2021) |
| $\boldsymbol{\theta_c}$ | Relative risk of SARS-CoV-2 transmission due to lockdown implementation | 0.23 | (Salje et al., 2020) |
| **Pneumococcal colonization and invasion parameters:** | | | |
| $\boldsymbol{\beta_S}$ | transmission rate of antibiotic-sensitive strain | 0.056 $days^{-1}$<br>0.046 $days^{-1}$<br>0.034 $days^{-1}$ | (Davies et al., 2019; Olesen et al., 2020) |
| $f$ | fitness of antibiotic-resistant strain (assuming there is a fitness cost on transmissibility) | 0.9652<br>0.949<br>0.926 | (Dagan et al., 2008; Melnyk et al., 2015) |
| $\boldsymbol{\beta_S f}$ | transmission rate of antibiotic-resistant strain | $\beta_S f$ | calculated |
| $\boldsymbol{\theta_\beta}$ | Relative risk of pneumococcal transmission due to lockdown implementation | 1 or 0.75 | assumed |
| $\boldsymbol{\gamma^S = \gamma^R = \gamma^{SS} = \gamma^{RR} = \gamma^{SR}}$ | rate of natural bacterial clearance (assumed to be the same for antibiotic-sensitive and -resistant strains) | 1/20 $days^{-1}$<br>1/30 $days^{-1}$<br>1/45 $days^{-1}$ | (Abdullahi et al., 2012; Davies et al., 2019; Ekdahl et al., 1997; Högberg et al., 2021; Melegaro et al., 2004) |
| $\boldsymbol{q}$ | relative infectiousness with each strain for dually colonized | 0.5 | assumed (Colijn et al., 2010) |
| $\boldsymbol{c}$ | fraction of dually colonized returning to single-colonized upon reinfection | 0.5 | assumed (Colijn et al., 2010) |
| $\boldsymbol{k}$ | probability of acquiring secondary bacterial carriage | 0.5 | assumed (Colijn et al., 2010) |
| $\boldsymbol{p_s}$ | probability of transmitting antibiotic-sensitive strain | 0.5 | assumed (Colijn et al., 2010) |
| $\boldsymbol{p_{single}}$ | probability of a single infection | 0.5 | assumed (Colijn et al., 2010) |
| $\boldsymbol{p_{inv}}$ | pneumococcal invasion rate (summer and winter) | [3x$10^{-6}$ $day^{-1}$, 9x$10^{-6}$ $day^{-1}$] in the elderly and general | (Domenech De Cellès et al., 2019; Opatowski et al., 2013) |

**Appendix 2**

| | | | |
|---|---|---|---|
| | | population, and [1x10$^{-6}$ day$^{-1}$, 2.5x10$^{-6}$ day$^{-1}$] in <5 years-old | |
| $S^S + S^R$ | initial states – initial prevalence of the total pneumococcal carriage (antibiotic-sensitive and -resistant) in different populations | 10%<br>20%<br>30% | (Cohen et al., 2023; Rose et al., 2021; Rybak et al., 2022; Tinggaard et al., 2023; Wang et al., 2017) |
| **Antibiotic exposure parameters:** | | | |
| $\omega$ | rate of antibiotic-induced pneumococcal clearance for sensitive strains (1/time before antibiotic action) | 1/3 days$^{-1}$ | (Kuitunen et al., 2023) |
| $r$ | rate of return to antibiotic unexposed compartment (1/duration of antibiotic treatment) | 1/7 days$^{-1}$ | (Grant and Saux, 2021; Kuitunen et al., 2023) |
| $r_{az}$ | rate of return to antibiotic unexposed compartment (1/the remainder of how long azithromycin stays in the body) | 1/11.5 days$^{-1}$ | calculated (Foulds et al., 1990; Girard et al., 2005) |
| $\tau$ | baseline rate of antibiotic exposure in the community (France) | 0.0014 average daily ppc (prescriptions per capita) | (Bara et al., 2022) |
| $a$ | A reduction factor for antibiotic exposure in the community resulting from changes in healthcare-seeking behavior in response to the COVID-19 pandemic | [0.51, 0.77, 0.84] to represent annual 13%, 18%, and 39% decrease observed in France | (Bara et al., 2022) |
| $p_{az}$ | A proportion of COVID-19 infected individuals in the community receiving azithromycin | [0-0.20] testing between 0% and 20% | (Tsay et al., 2022; Wittman et al., 2023) |
| **Pathogenicity (invasive pneumococcal disease risk):** | | | |
| $IPD_{risk}$ | A reduction factor for the risk of developing an invasive pneumococcal disease (IPD) due to the absence of influenza-like-illnesses (ILIs) after lockdown implementation | 1 (pre-lockdown)<br>0.2 (lockdown)<br>0.4 (post-lockdown)<br>for an average of 0.5 in 2020 | (Shaw et al., 2023) |
